# mTORC2-mediated cell-cell interactions promote BMP4-induced WNT activation and mesoderm differentiation

**DOI:** 10.1101/2024.06.07.597881

**Authors:** Li Tong, Faiza Batool, Yueh-Ho Chiu, Priscilla Di Wu, Yudong Zhou, Xiaolun Ma, Santosh Atanur, Wei Cui

## Abstract

The mechanistic target of rapamycin complex 2 (mTORC2) is essential for embryonic development but its underlying molecular mechanisms remain unclear. Here we show that disruption of mTORC2 in human embryonic stem cells (hESCs) considerably alters the Rho/Rac signaling dynamics and reduces cell adhesion. Despite this, mTORC2-deficient hESCs maintain self-renewal and expression of pluripotent markers when cultured in mouse-embryonic fibroblast conditioned medium supplemented with bFGF (MEF-CM). However, these hESCs exhibit significantly impaired mesoderm and endoderm differentiation in response to BMP4 and Activin, respectively, due to reduced WNT activation mediated by cell-cell interactions. Direct activation of the WNT pathway by a GSK3 inhibitor restores mesendoderm differentiation in mTORC2-deficient hESCs. Our study uncovers a novel mechanism by which mTORC2 regulates cell fate determination and highlights a critical link between the intercellular adhesion and the activation of canonical WNT genes.

## Introduction

The mechanistic/mammalian target of rapamycin (mTOR) is a serine/threonine protein kinase that is evolutionarily conserved in all eukaryotes. It functions through two distinct multiprotein complexes, mTORC1 and mTORC2, both of which form homodimers (Aylett et al., 2016; Scaiola et al., 2020). Although both complexes contain mTOR as the catalytic subunits and mammalian lethal with SEC13 protein 8 (mLST8), each complex also contains specific components. RAPTOR (regulatory-associated protein of mTOR), which is specific to mTORC1, is replaced by RICTOR (rapamycin-insensitive companion of mTOR) and mSIN1 (mammalian stress-activated map kinase-interacting protein 1) in mTORC2. These complex-specific components dictate their distinct regulatory mechanisms, substrates, and functions (Figure 1A) (Battaglioni et al., 2022). mTOR signaling plays an important role in cell proliferation, cytoskeleton remodeling and cell metabolism; hence, it is essential for embryonic development and tissue homeostasis (Liu and Sabatini 2020). Over the past decades, significant advances have been made in understanding of mTORC1 (Battaglioni et al., 2022; Kim and Guan 2019), while the functions and underlying molecular mechanisms of mTORC2 remain less well-defined due to its intricate crosstalk with other pathways, interwoven regulatory machinery and the absence of mTORC2-specific inhibitors.

**Figure 1.**
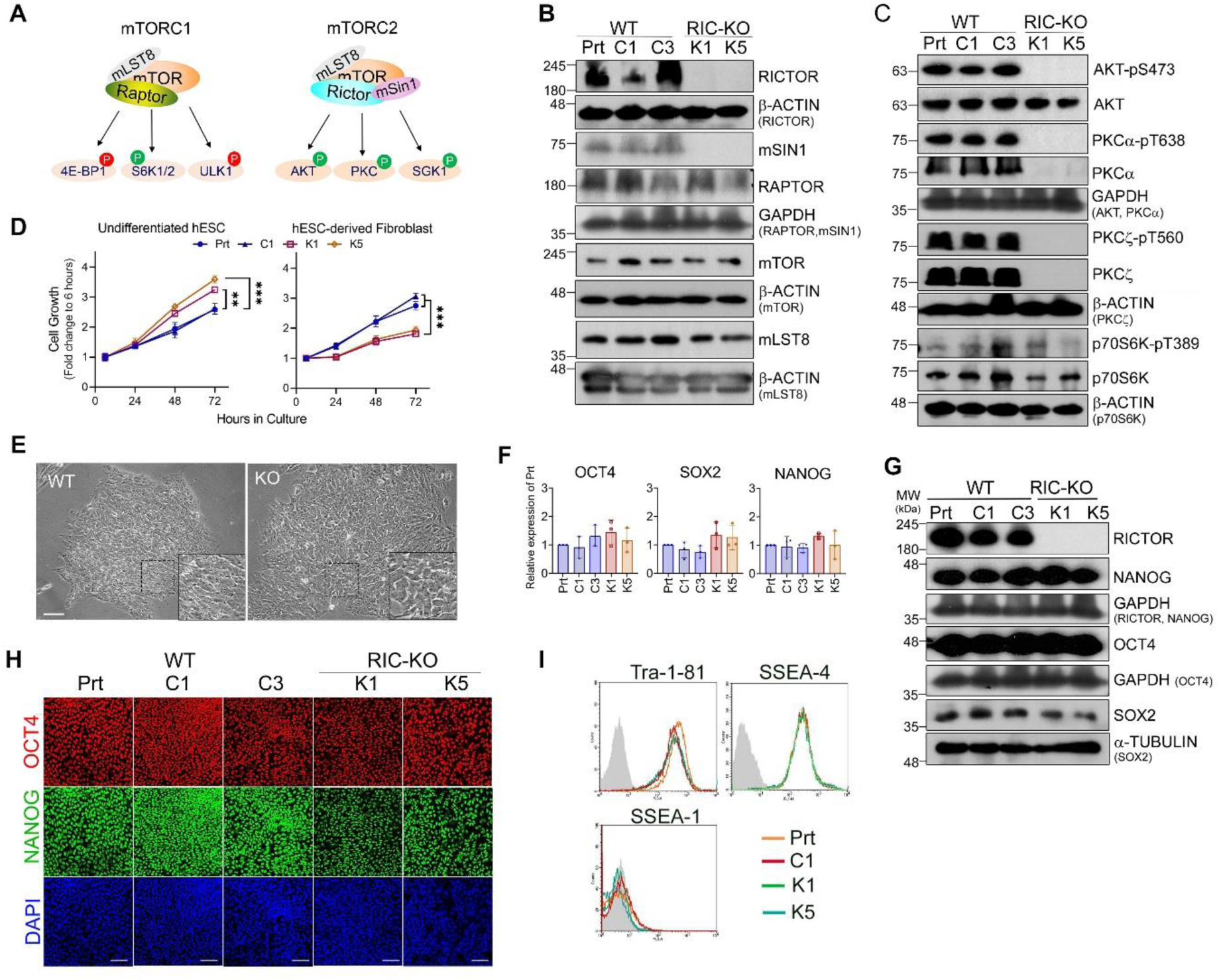
RICTOR-knockout hESCs retain the self-renewal and pluripotent marker expression. (A) Schematics illustrate mTOR complexes their promoting and inhibitory substrates, green and red P, respectively. (B,C) The expression of mTORC1 and mTORC2 key subunits (B) and the phosphorylation on their targets (C) (n > 2) by immunoblotting. Prt, parental H1 hESCs; C1 & C3, WT controls; K1 & K5, RIC-KO clones. (D) Proliferation of hESC lines and their differentiated fibroblasts by CCK8 assay. Data are presented as mean ± SEM (n=3). ** & ***, *p* <0.005 & 0.0005, respectively by two-way ANOVA. (E) Phase-contrast images of WT and RIC-KO hESCs. Scale bar = 50 µm. (F) Expression of key pluripotent transcription factors in hESCs by RT-qPCR. Data are presented as mean ± SD (n = 3). (G,H) Protein expression of key pluripotent transcription factors in hESCs by immunoblotting (G) and immunostaining (H) (n > 2). Scale bar = 25 µm. (I) Expression of hESC-associated cell surface antigens in hESCs by flow cytometry (n = 2). See also Figure S1.

Loss-of-function studies in mice demonstrate that mTORC2 is vital for embryonic development. Disruption of either *Rictor* or *mSin1* leads to embryonic lethality at E10-11 (Guertin et al., 2006; Shiota et al., 2006), which coours later than the death of Raptor-null embryos at E6.5, suggesting that mTORC1 and mTORC2 have diverse functions in embryonic development. Intriguingly, tissue-specific knockout of mTORC2 in mice, such as in endothelial cells, generally results in mild phenotypes without overt developmental defects, unless the disruption occurs around gastrulation; in such cases, the phenotype closely resembles that of a germline knockout (Aimi et al., 2015; Bentzinger et al., 2008; Cybulski et al., 2009; Dragert et al., 2015; Godel et al., 2011; Hagiwara et al., 2012; Lamming et al., 2012; Zhao et al., 2014). These imply that mTORC2 may have an important function during early embryonic development, particularly around gastrulation, yet the underlying molecular mechanisms are largely unknown. Additionally, the role of mTOR signaling, especially mTORC2, in pluripotent stem cells (PSCs) also remains elusive (Bulut-Karslioglu et al., 2016; Zhou et al., 2009).

To explore the role of mTORC2 in embryonic development and PSCs, we disrupted the expression of *RICTOR* in human embryonic stem cells (hESCs) and characterized the resulting cells for their self-renewal and cell fate determination. *RICTOR* knockout (RIC-KO) hESCs exhibit a considerable reduction in cell adhesion and interactions with a decreased propensity for mesoderm and endoderm differentiation. Further studies revealed that mTORC2-mediated cell interactions are crucial for mesoderm and endoderm differentiation through modulation of WNT signaling.

## Results

### Eradicating mTORC2 signaling shows no effect on self-renewal of hESCs

The *RICTOR* gene was disrupted in H1 hESCs using CRISPR/Cas9 and two RIC-KO clones (K1 and K5) were expanded in MEF-CM (Figure S1A,B). Elimination of *RICTOR* dramatically reduced mSIN1 protein levels without affecting other core subunits of the mTOR complexes as reported before (Yang et al., 2006) (Figure 1B). As expected, mTORC2 activity was abolished, while mTORC1 signaling remained (Figure 1C). During initial clonal expansion, RIC-KO hESCs showed a transient reduction in cell growth which gradually diminished. When clonal lines were established, RIC-KO hESCs revealed faster growth than WT controls with increased MAPK-ERK pathway activity (Figure 1D; Figure S1C). Interestingly, upon differentiation into fibroblasts, these RIC-KO cells grew slower than WT cells (Figure 1D), which is in line with fibroblasts derived from *Rictor*-null mouse embryos (Guertin et al., 2006; Shiota et al., 2006). Both RIC-KO hESC lines maintained normal chromosome numbers and formed alkaline phosphatase-positive colonies similar to WT controls (Figure S1D,E; Figure 1E). Expression of key pluripotent transcription factors and hESC-associated cell surface antigens showed no clear change (Figure 1F-I). However, when cultured in chemically defined mTeSR medium, the RIC-KO hESCs gradually lost their pluripotent features (Figure S1F,G) as previously reported (Chu et al., 2022). These results suggest that culture conditions have crucial effects on RIC-KO hESCs.

To circumvent off-target effects of sgRNAs and cell line-dependent bias, *RICTOR* expression was also knocked down in H7 hESCs using shRNA (RIC-KD) (Figure S1H,I). These cells exhibited similar growth dynamics, morphology, and marker expression to H1 RIC-KO hESCs (Figure S1J-L). Together, these data demonstrate that disrupting RICTOR in hESCs eliminates mTORC2 activity without affecting mTORC1 signaling and that mTORC2 is not essential for the maintenance of hESC self-renewal in the MEF-CM culture conditions.

### mTORC2-deficiency in hESCs significantly reduces cell adhesion and alters RhoA/Rac signaling

In our routine culture, hESCs were propagated by collagenase-assisted mechanical splitting and small cell clumps were plated at a 1:3 ratio onto Matrigel-coated plates (Noisa et al., 2011). Although *RICTOR* disruption showed no apparent effect on hESC self-renewal, both H1 RIC-KO and H7 RIC-KD (together called RICTOR-deficient, RIC-DF) hESCs exhibited noticeably reduced attachments to culture plates, but this defect was visibly ameliorated by the addition of the ROCK inhibitor Y-27632 (ROCKi) (Figure 2A,B; Figure S2A). Furthermore, intercellular junction proteins, including E-cadherin (E-CAD) and ZO1, showed lower levels in RIC-DF hESCs without significant changes on the transcript levels (Figure 2C,D; Figure S2B-D). β-CATENIN (β-CAT) protein also displayed reduced localization at the cell membrane. These findings are consistent with the observation that RIC-DF hESC colonies were less compact than WT controls (Figure 1E; Figure S1K), suggesting reduced cell interactions in RIC-DF hESCs.

**Figure 2.**
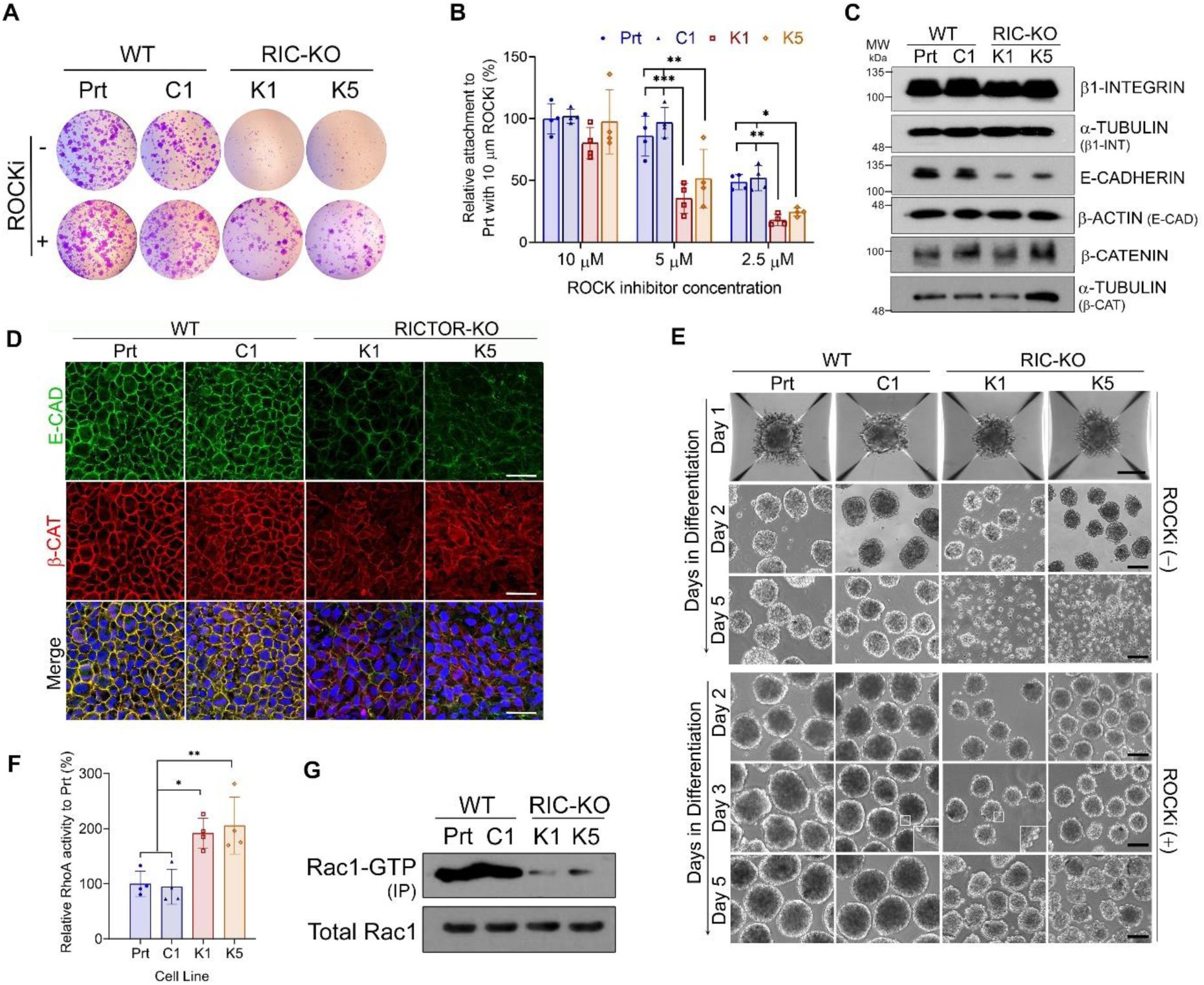
RIC-KO hESCs exhibit reduced cell adhesion. (A) Images of violet staining of hESC colonies 24 hours after plating in the presence or absence of the ROCK inhibitor (ROCKi) (n = 3). (B) Cell attachment assay with indicated concentration of ROCKi. Data are presented as mean ± SD (n = 4). *, ** & ***, *p* < 0.05, 0.01 and 0.0005 by two-way ANOVA. (C) Expression of proteins associated with cell adhesion by immunoblotting (n ≥ 2). (D) Immunostaining of WT and RIC-KO hESCs with indicated antibodies. Scale bar = 30 µm. (E) Phase-contrast images of EB formation (n = 2) in hESCs in the absence or presence of ROCKi (n = 2). Inserts on day 3 showing blebs in RIC-KO EBs. Scale bar = 100 µm. (F) RhoA activity in hESCs. Data are presented as mean ± SD (n = 4). * & **, *p* < 0.05 and 0.005, respectively, by one-way ANOVA. (G) Rac1-GTP pull down assay in hESCs (n = 2). See also Figure S2.

The impeded cell interactions in RIC-DF hESCs became more evident during embryoid body (EB) formation. In contrast to WT hESCs that require ROCKi only on the first day (Figure S2E), RIC-DF hESCs required continuous ROCKi treatment throughout EB formation; without it, cell aggregates disintegrated (Figure 2E). Notably, the disintegrated cells remained viable, as indicated by negative trypan blue staining. Even with ROCKi, RIC-DF EBs showed more surface blebbing (Figure 2E; Figure S2F). These results reveal that RICTOR/mTORC2-deficiency significantly hinders cell-ECM (extracellular matrix) and cell-cell interactions.

Since ROCKi partially rescued the attachment defects of RIC-DF hESCs, RhoA-ROCK signaling was thought to be higher in these cells. Indeed, RhoA activity was significantly increased in RIC-DF hESCs (Figure 2F; Figure S2G), while the levels of Rac1-GTP were reduced (Figure 2G; Figure S2H). Phalloidin staining further showed an altered pattern of F-ACTIN organization in RIC-KO hESCs, implying a change in cytoskeleton (Figure S2B). RhoA and Rac1 are well known to function antagonistically and coordinately regulate cell adhesion, cytoskeleton, and movement (Burridge and Wennerberg 2004; Martin et al., 2016; Yang et al., 2006). Our results suggest that mTORC2 plays an important role in maintaining the balance of RhoA-ROCK/Rac1 signaling in hESCs and that disruption of mTORC2 considerably upregulates RhoA-ROCK, contributing to impaired cell adhesion and cytoskeletal disorganization.

### Disrupting mTORC2 in hESCs downregulates genes involved in mesoderm differentiation and WNT signaling

To assess the impact of mTORC2 disruption in hESCs, RNA-seq analysis was performed on WT (C1 & C3) and RIC-KO (K1 &K5) hESCs. The vast majority of genes showed no significant change in expression and only 0.6% (195 out of 32673 genes) were differentially expressed (> 2-fold change, adjusted *p* < 0.05), including 140 downregulated and 55 upregulated genes (Figure 3A,B; Figure S3A; also see Table S1,S2). Notably, several downregulated genes encode transcription factors vital for lineage specification, particularly mesoderm and endoderm, including *GATA4*, *FOXA2, SOX17 and SP5* (Mukherjee et al., 2020; Rojas et al., 2005; Weidinger et al., 2005). In contrast, genes associated with pluripotency or neural differentiation were expressed at the similar levels (Figure 3B; Figure S3B). Since mTORC2 primarily regulates protein phosphorylation, it is not surprising that genes directly involved in cell adhesion, such as E-cadherin, were not transcriptionally altered (Figure S2C). Although no statistically significant results were obtained on the upregulated genes by Gene ontology (GO) enrichment analysis, the analysis of the downregulated genes showed significant association with embryonic development, embryo patterning, and muscle formation, as well as regulation of gene expression and growth factor/cytokine activity (Figure 3C,D; Table S3). KEGG pathway analysis further linked these genes to WNT signaling (Figure 3E). These findings suggest that mTORC2 may have a role in hESC differentiation by modulating the expression of lineage-specific genes, possibly through regulation of WNT signaling.

**Figure 3.**
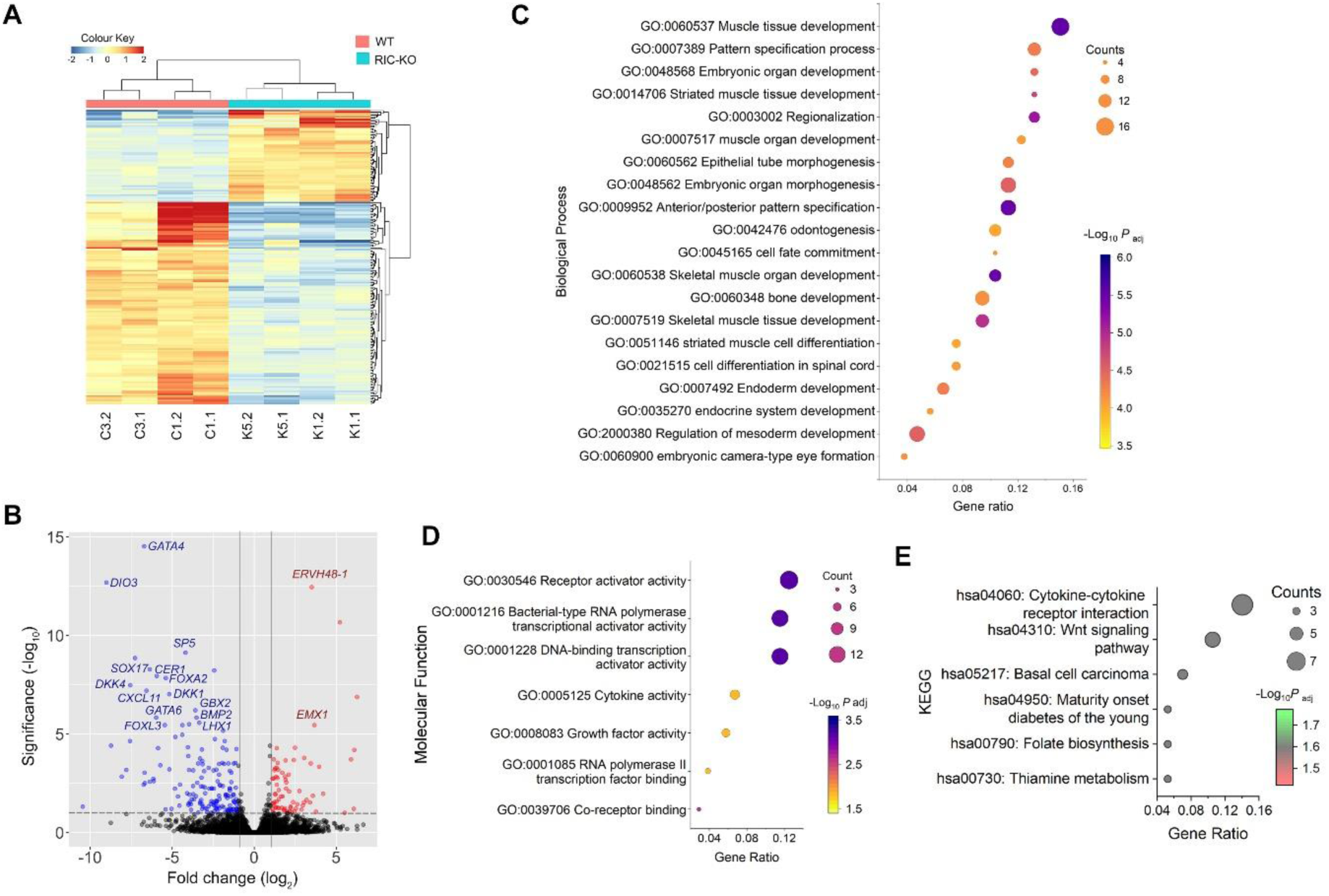
Altered expression of developmental-associated genes in RIC-KO hESCs by RNA-seq. (A) Heatmap and hierarchical clustering showing differentially expressed genes in WT and RIC-KO hESCs (n = 2). (B) Volcano plots of differentially expressed genes (adjusted *P* value (*Padj*) < 0.05; log2 (fold change) > 1) in RIC-KO hESCs. Blue and red dots represent downregulated and upregulated genes, respectively. (C,D) Gene ontology enrichment of downregulated genes in biological process (C) and molecular Function (D) (*Padj* < 0.05). (E) KEGG pathway enrichment (*Padj* < 0.05) of downregulated genes. See also Figure S3.

### mTORC2-deficiency hinders mesoderm and endoderm differentiation of hESCs

To evaluate the differentiation potential of RIC-DF hESCs, EB-mediated differentiation was performed in the presence of ROCKi (Figure S2E). Although RIC-DF EBs contained derivatives of all three germ layers, the proportions of mesodermal and endodermal cells were much less than WT controls (Figure 4A, Figure S4A). This was corroborated by mRNA expression analysis, showing significantly lower levels of mesoderm (*BRACHYURY*, *TBX6* and *FLK1*) and endoderm (*FOXA2*, *GATA6* and *SOX17*) markers in RIC-DF EBs (Figure 4B, Figure S4B).

**Figure 4.**
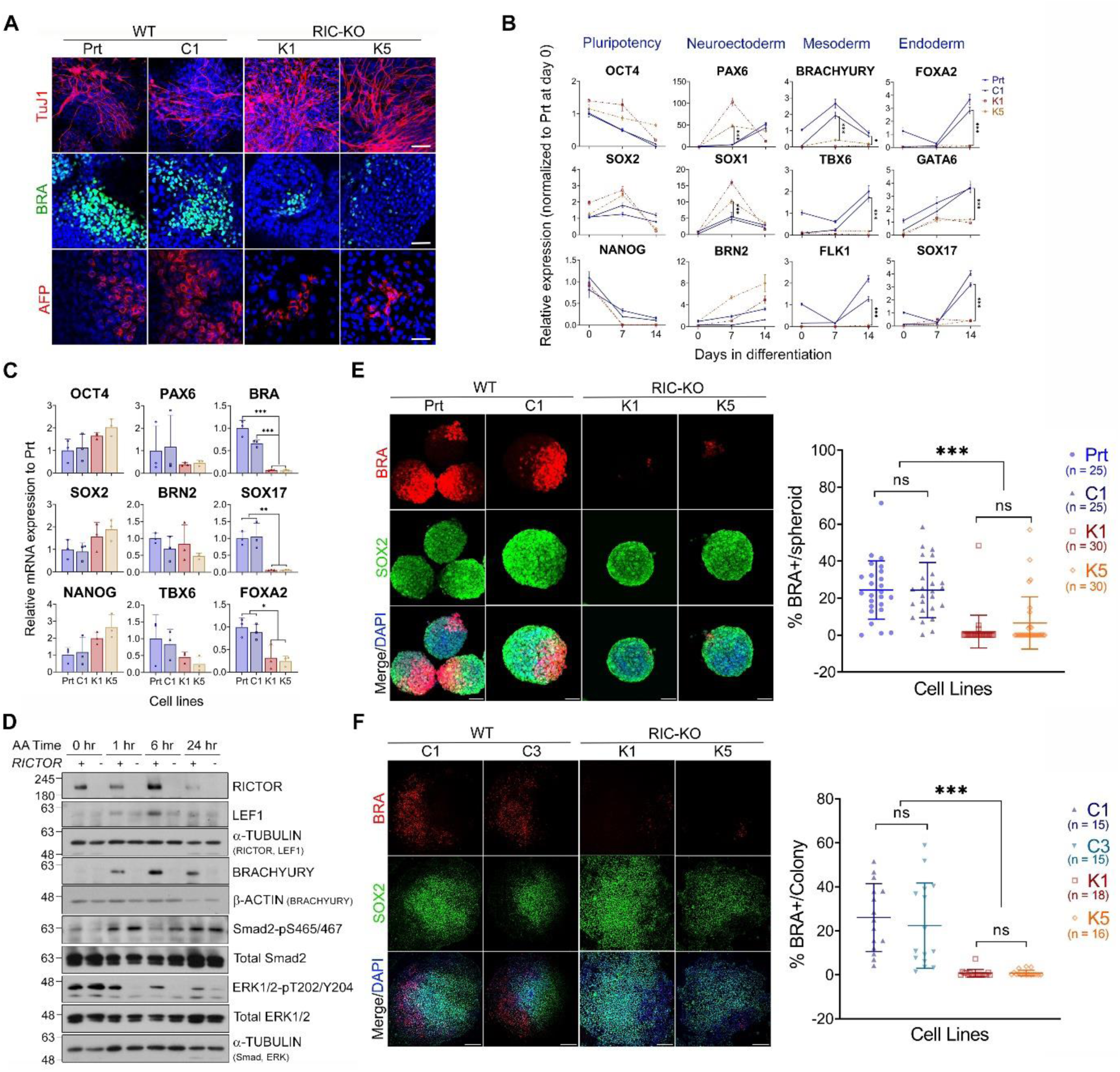
mTORC2-deficiency impedes mesoderm and endoderm differentiation. (A) Immunostaining of the three germ layer markers after 14-day differentiation via EB (n = 2). Scale bar = 50 µm. (B) Dynamic mRNA expression during EB differentiation by RT-qPCR. Data are presented as mean ± SD of 6 measurements of 2 independent differentiation experiments. (C) Expression of indicated pluripotent and lineage genes after 72 hours of Activin-induced endoderm differentiation. Data are presented as mean ± SD (n = 3). (D) Expression of indicated proteins during Activin-induced endoderm differentiation by immunoblotting in WT and RIC-KO hESCs. (E) Max intensive projection images of immunostaining with Brachyury (BRA) and SOX2 antibodies (left, scale bar = 75 µm) and quantification of % BRA+ cells/spheroid (right). (F) Images of immunostaining with Brachyury (BRA) and SOX2 antibodies (left, scale bar = 250 µm) and quantitative analysis (right). The quantification in E & F is presented as mean ± SD with each symbol representing a spheroid/colony with the total number analyzed of each line from 3 independent experiments indicated in the bracket. *, ** & ***, *p* < 0.05, 0.005, and 0.0005 by one-way ANOVA. See also Figure S4.

To further assess endodermal differentiation, RIC-KO hESCs were also subjected to Activin A (AA)-induced differentiation (Figure S4C), a process previously shown to be negatively regulated by PI3K/mTORC2 signaling (Yu et al., 2015). Unexpectedly, RIC-KO hESCs exhibited significantly impaired endoderm differentiation despite the prolonged AA-induced pSmad2/3 signal duration (Figure 4C,D), consistent with prior findings (Yu et al., 2015). Notably, expression of BRACHURY, a mesendoderm marker, was also reduced in RIC-KO cells (Figure 4C,D), indicating that impaired mesendoderm might underlie the observed deficiency in endoderm differentiation. This aligns with the role of mesendoderm as a bipotent precursor for both mesoderm and endoderm, resembling cells of the primitive streak in gastrulating embryos (Rodaway and Patient 2001).

To further validate the role of mTORC2 in mesendoderm differentiation, we adapted a BMP4-induced differentiation protocol in 2D-colony and 3D-spheroid cultures (Figure S4D,E) (Bernardo et al., 2011; Simunovic et al., 2019). After 48-hours of BMP4 induction, WT hESCs exhibited symmetry breaking, characterized by the emergence of BRACHYURY-positive (BRA+) cells one side of a colony or spheroid and SOX2-positive cells on the opposite side (Figure 4E,F), consistent with previous results (Simunovic et al., 2019). In contrast, RIC-KO cultures hardly showed BRA+ cells, indicating a failure to initiate mesendoderm differentiation. Correspondingly, expression of mesoderm/endoderm genes was significantly reduced in RIC-KO cells (Figure S4F). Taken together, these results demonstrate that ablation of mTORC2 in hESCs considerably impedes their mesendoderm differentiation, implying that mTORC2 has a regulatory role in early lineage specification.

### Activation of WNT signaling recovers mesendoderm differentiation in RIC-KO hESCs

Next, to investigate the underlying mechanisms by which mTORC2 regulates mesendoderm differentiation, we focused on BMP, Wnt and Nodal signaling pathways which are known to orchestrate gastrulation in mouse embryos (Nowotschin and Hadjantonakis 2020). In this signaling cascade, BMP4 secreted from extraembryonic tissues activates Wnt signaling in epiblast, which in turn enhances Nodal, collectively promoting the formation of the primitive streak/mesendoderm (Figure S5A) Similar regulatory mechanisms have been observed in non-human primates, indicating their conservation between species (Cui et al., 2022). To test whether mTORC2-deficient hESCs are competent to respond to these signals, both WT and RIC-KO hESCs were treated with BMP4, CHIR99021 (CHIR; a GSK3 inhibitor to activate WNT signaling), and AA (a NODAL pathway agonist) (Figure S5B). This combined treatment substantially increased the proportion of BRA+ cells in RIC-KO cultures, even exceeding the proportion observed in WT controls (Figure 5A). These results suggest that the RIC-KO hESCs are competent to differentiate into mesendoderm and that mTORC2-deficiency possibly disrupts BMP4-induced activation of WNT and/or NODAL pathways. To dissect this further, both hESCs were treated with BMP4 in combination with either CHIR or AA. While AA did not improve BMP4-induced BRA+ cells in RIC-KO cells (Figure 5B), the addition of CHIR dramatically increased BRA+ cells and eliminated the differences between WT and RIC-KO cultures (Figure 5C). Moreover, CHIR alone was sufficient to induce high levels of BRA⁺ cells in both cultures (Figure 5D). These results imply that mTORC2-deficiency impedes mesendoderm differentiation by encumbering BMP4-induced activation of WNT signaling which is essential for mesendoderm differentiation in hESCs.

**Figure 5.**
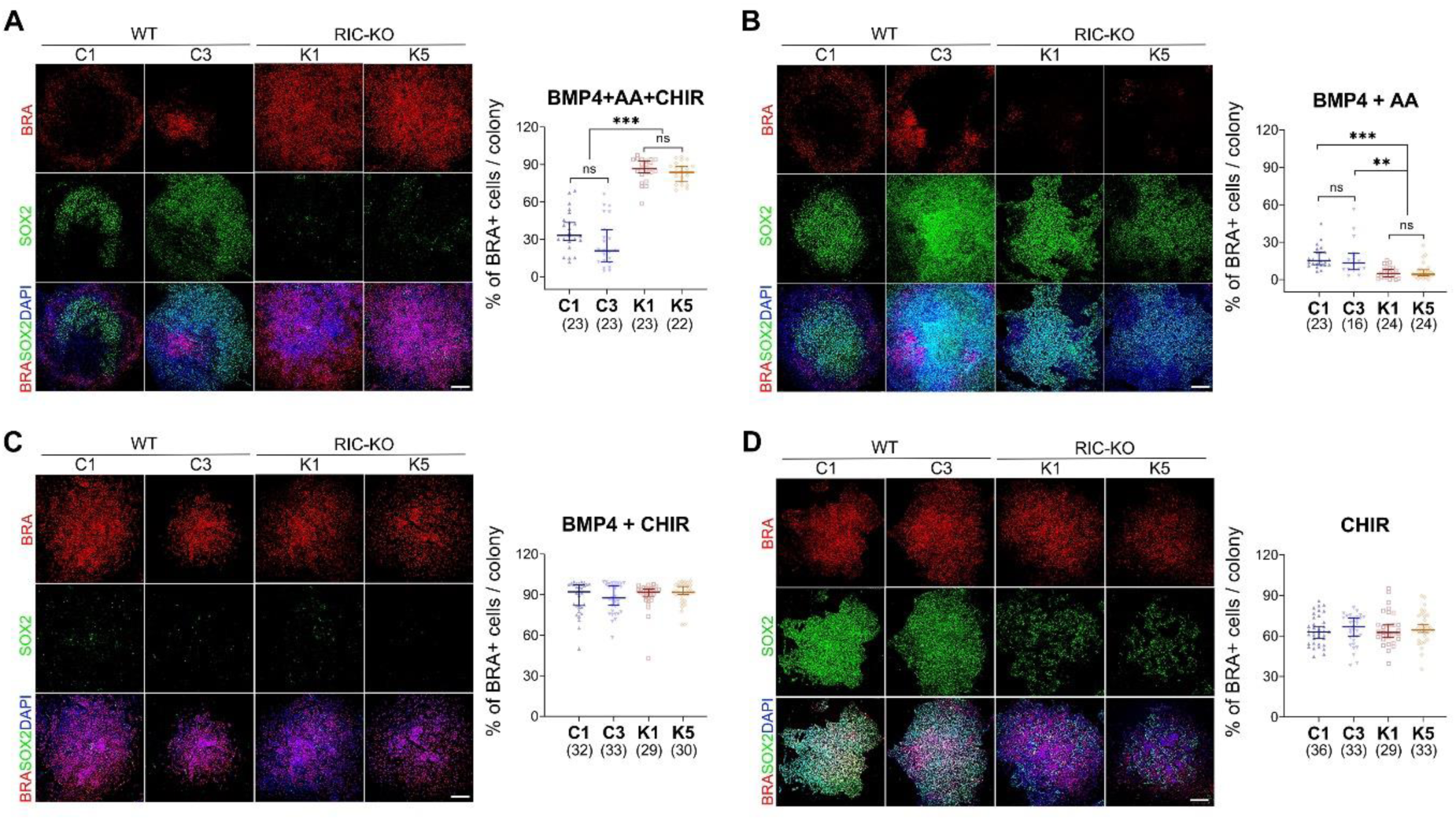
Activation of WNT considerably improves mesendoderm differentiation in RIC-KO hESCs. Immunostaining images of Brachyury (BRA) and SOX2 (left, Scale bar = 250 µm) after 48 hours of the indicated treatment. Quantification of BRA+ cells (right) is presented as median with 95% confidence interval. Each symbol represents a colony. The total number of colonies analyzed in each line is shown in brackets (n = 3). ** & ***, *p* < 0.005 and 0.0005, respectively, by one-way ANOVA. (A) BMP4 with Activin A (AA) and CHIR99021 (CHIR); (B) BMP4 and AA; (C) BMP4 and CHIR; (D) CHIR only. See also Figure S5.

Interestingly, when comparing the effects of these treatments within the same cell lines, we observed that AA combined with BMP4 did not significantly increased BRA+ cells compared to BMP4 alone (BMP4+AA vs BMP4). However, in WT hESCs, the addition of AA to BMP4 and CHIR (BMP4+AA+CHIR) reduced the proportion of BRA+ cells compared to BMP4 + CHIR, which was not observed in RIC-KO cells (Figure S5C). The underlying mechanisms behind the context-dependent modulation remain to be elucidated. Together, these results further support the notion that mTORC2 regulates BMP4-induced mesendoderm differentiation primarily through modulating the activation of WNT signaling.

### BMP4-induced expression of WNT genes is impeded in RIC-KO hESCs

Given that GSK3 inhibition efficiently induced mesendoderm differentiation in RIC-KO hESCs (Figure 5), we expected that mTORC2 may affect the WNT pathway upstream of GSK3. Indeed, BMP4 significantly upregulated canonical WNT genes, *WNT3, WNT3A* and *WNT8A*, in WT, but not in RIC-KO hESCs, and the expression of WNT target genes, *AXIN2* and *LEF1*, was also higher in WT cells (Figure 6A,B). Although mTORC2 is known to maximize AKT activity which may exert inhibitory phosphorylation on GSK3 and thereby activate WNT signaling (Nusse and Clevers 2017), no clear difference was detected on the levels of GSK3 phosphorylation between WT and RIC-KO cells (Figure 6B; Figure S6A). Therefore, mTORC2-deficiency appears to suppress BMP4-induced canonical WNT activation, thereby reducing WNT signaling and impairing mesendoderm differentiation in hESCs.

**Figure 6.**
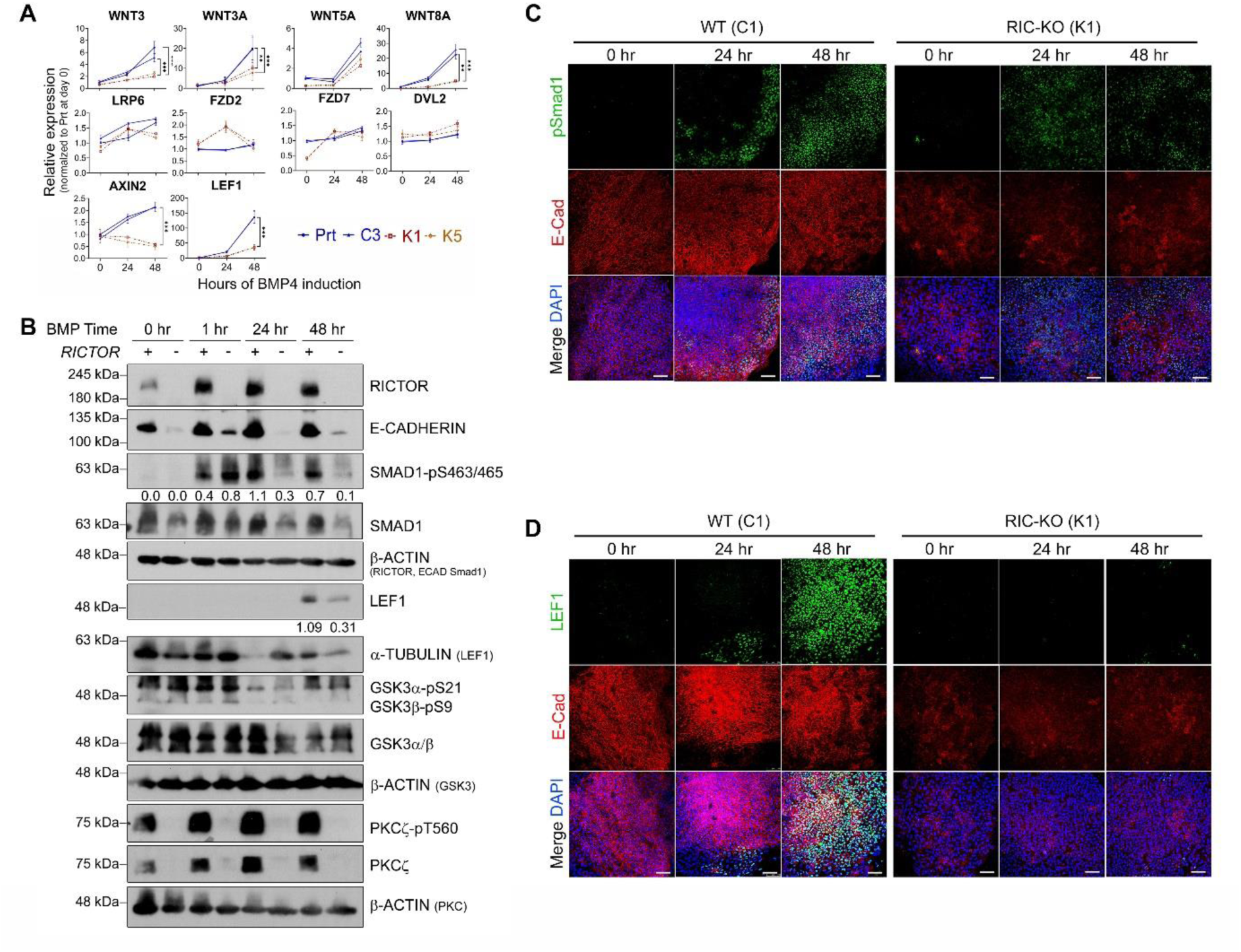
BMP4-induced activation of WNT is hampered in RIC-KO hESCs. (A) Expression of genes associated with WNT signaling during BMP4-induced differentiation by RT-qPCR. Data are presented as mean ± SD (n = 3). ** & ***, *p* < 0.005 and 0.0005, respectively, by two-way ANOVA. (B) Immunoblots showing protein expression in WT and RIC-KO hESCs at the indicated timepoints during BMP4-induced differentiation (n = 2). Numbers below the pSmad1 and LEF1 bands indicate the relative signal intensity normalized to the loading control using ImageJ. (C,D) Immunostaining with the indicated antibodies at the specified timepoints during BMP4- induced differentiation in C1 and K1 hESCs. Scale bar = 100 µm. (n = 3). See also Figure S6.

BMP4 receptors have been reported to be localized at the basolateral sides of cells, meaning that only the peripheral cells of a colony are accessible to externally applied BMP4, and E-CAD has been implicated in modulating WNT signaling patterning in hESCs (Etoc et al., 2016; Martyn et al., 2019; Zhang et al., 2019). Since RIC-KO hESCs exhibited a marked reduction in E-CAD expression and cell interactions (Figure 2), we asked whether these changes could affect BMP4 and WNT signaling. Although both WT and RIC-KO hESCs showed similar acute responses to BMP4, phosphorylation of SMAD1/5/9 (pSMAD1) dramatically declined at 24 hours in RIC-KO cells only (Figure 6B). This could be due to higher upregulation of the BMP4 antagonist NOGGIN in RIC-KO hESCs (Figure S6B). Furthermore, the emergence of pSMAD1 signals in WT coincided with the loss of E-CAD within WT colonies (Figure 6C; Figure S6C), and more importantly, LEF1 exhibited a spatial pattern similar to that of pSMAD1 (Figure 6D; Fig. S6D). Collectively, these findings suggest that cell-cell interactions may contribute not only to the BMP4 signaling gradient but also to the activation of WNT signaling.

### Elimination of cell interactions diminishes BMP4-induced upregulation of WNT genes

To further validate the role of cell-cell interactions in activating WNT genes and mesendoderm differentiation in hESCs, we performed the differentiation using WT hESCs cultured either in colony (CL) or single cell (SC) form (Figure 7A). As expected, asymmetrically distributed BRA+ cells were detected in CL cultures after 48 hours of BMP4 treatment, while BRA+ cells were almost absent in SC cultures (Figure 7B). In SC cultures, pSMAD1 signals were diffuse, and membrane-bound E-CAD was undetectable (Figure 7C). Expression of canonical WNT ligands, WNT target genes, and mesendoderm markers was lower in SC than in CL cultures, (Figure 7D; Figure S7A). However, simultaneous activation of BMP, WNT and AA pathways in the SC cultures efficiently induced BRA+ cells (Figure S7B). Together, These results support the finding that BMP4-induced mesendoderm differentiation in hESCs requires close cell-cell contact as such interactions are vital for the activation of WNT genes.

**Figure 7.**
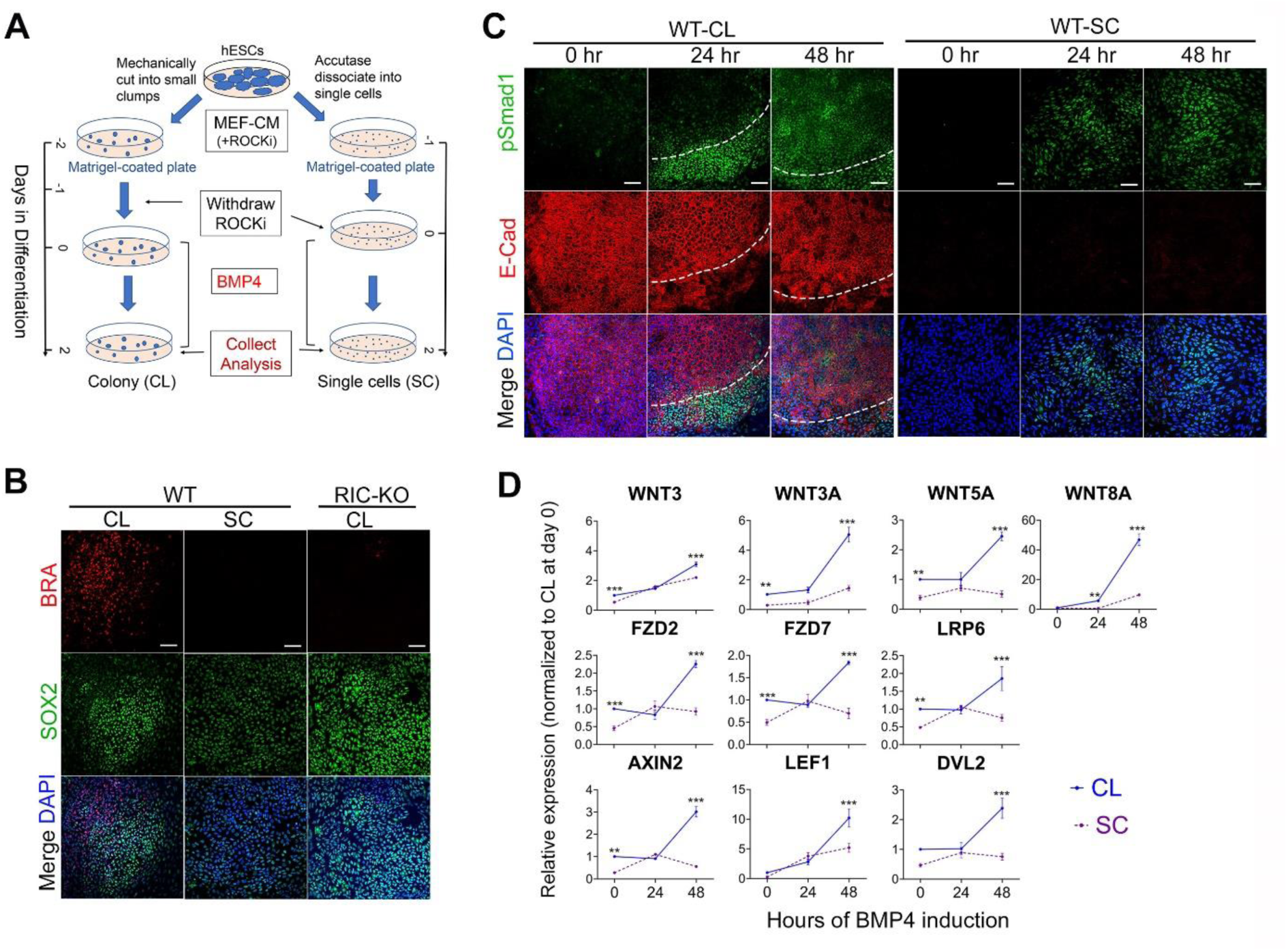
Cell interactions promote BMP4-induced activation of WNT in hESCs. (A) Schematic of the differentiation procedure in WT hESCs as colonies (CL) and single cells (SC). (B) Immunostaining for BRA and SOX2 after 48 hours of BMP4 treatment in CL and SC cultures. Scale bar = 100 µm. (C) Immunostaining for pSMAD1and E-CAD at the indicated timepoints during BMP4- induced differentiation in CL and SC cultures (n = 3). (D) Expression of genes associated with WNT signaling during BMP4-induced differentiation in CL and SC cultures by RT-qPCR. Data are presented as mean ± SD (n = 3). ** & ***, *p* < 0.005 and 0.0005, respectively, by two-way ANOVA. See also Figure S7.

## Discussion

In this study, we investigated the role of mTORC2 in regulating PSC properties using RICTOR-deficient hESCs. We show that although RICTOR disruption initially causes a transient proliferative reduction, the hESCs adapt over time and exhibit enhanced growth without clear genetic alterations. This suggests a capacity for adaptive signaling rewiring, potentially involving upregulation of FGF-MAPK-ERK pathway. Interestingly, this growth recovery is observed only in undifferentiated hESCs while differentiated derivatives (e.g. fibroblast and EB) show growth inhibition, which indicates that the growth recovery reflects adaptive rewiring rather than transformation. Furthermore, our results show that the self-renewal of RIC-KO hESCs can only be maintained in MEF-CM but not in defined mTeSR conditions (Chu et al., 2022). It is possible that MEF-CM provides ECM and extracellular vesicles that compensate, at least partially, for impaired intercellular communication in mTORC2-deficient hESCs (Ahmed and Ffrench-Constant 2016; Kalluri and McAndrews 2023). Therefore, our results demonstrate that mTORC2 is not essential for the maintenance of hESCs.

Mechanistically, we identify that mTORC2 influences Rho family of GTPases which is crucial for cell adhesion and epithelial morphogenesis (Heasman and Ridley 2008; Jaffe and Hall 2005). In RIC-DF hESCs, increased RhoA-GTP and reduced Rac1-GTP lead to aberrant cell adhesion, which is partially rescued by ROCK inhibition. mTORC2 may regulate these GTPases through multiple pathways, such as PKC-mediated phosphorylation of RhoA/Rac regulators, including Rac1-specific guanine nucleotide exchange factor (GEF), RhoGDI2 and p190RhoGAP (Griner et al., 2013; Levay et al., 2009; Morrison et al., 2015). As a substrate of mTORC2, PKC is misfolded and degraded upon mTORC2 disruption (Ikenoue et al., 2008), which could affect RhoA/Rac1 signaling. Furthermore, RICTOR has been shown to suppress RhoGDI2 independent of mTORC2 activity to maintain Rac1 GTPase activity (Agarwal et al., 2013).

A main finding of our study is that mTORC2-deficiency considerably impairs mesoderm and endoderm differentiation. This impairment is not due to a loss of intrinsic lineage potential but rather to defective activation of canonical WNT signaling. Forced activation of the WNT/β-CATENIN pathway fully rescues mesendoderm differentiation in RIC-KO hESCs, underscoring the essential role of WNT signaling in this process (Steinhart and Angers 2018). Importantly, we demonstrate that cell-cell interactions are necessary for BMP-induced expression of canonical WNT ligands in hESCs. While the regulation of WNT signaling has been intensively studied, the research has focused on downstream components with limited attention to the regulation of upstream WNT ligands. Previous studies have shown that the p53 family directly activates the *Wnt3* gene in mouse ESCs to promote their mesendoderm differentiation (Wang et al., 2017) and that YAP protein can bind to an intragenic enhancer of the *WNT3* gene in hESCs to prevent its premature activation by Activin-induced SMAD2/3 (Estaras et al., 2017). Here we show that mTORC2-mediated cell interactions are vital for BMP4-induced canonical WNT gene activation and spatial patterning during early differentiation. These interactions may also affect Activin-Smad2/3-induced WNT activation as RIC-KO hESCs show reduced LEF1 to Activin. How cell-cell interactions regulates WNT gene activation and whether this is related to p53 or YAP in hESCs await further investigation.

In addition to WNT activation, mTORC2 also differentially modulates BMP and Activin pathways. Specifically, mTORC2 deficiency shortens BMP-Smad1/5 signaling duration but prolongs Activin-Smad2/3 signaling. These effects may be influenced by differences in culture media and the distinct linker sequences between Smad1/5 and Smad2/3 (Macias et al., 2015). Furthermore, mTORC2 deficiency affects MAPK-ERK signaling in a context-dependent manner. RIC-KO hESCs show increased FGF-ERK signaling in MEF-CM but reduced ERK signaling in Activin-induced endoderm cultures, potentially impacting pSmad2/3 signaling (Richardson et al., 2023). Interestingly, while transient mTOR inhibition promotes endoderm formation, complete mTORC2 knockout hinders it, despite sustained Activin-Smad2 signaling. This discrepancy may be explained by the requirement for cell interaction-mediated WNT activation, which is disrupted in RIC-KO cells. Collectively, these findings highlight the integral role of mTORC2 in coordinating multiple signaling pathways during hESC differentiation.

Mesendoderm differentiation recapitulates key aspects of primitive streak formation during gastrulation. Although mTORC2-deficient hESCs exhibit defects in mesendoderm differentiation, mTORC2-null mouse embryos can undergo successful gastrulation and form all three germ layers (Guertin et al., 2006; Shiota et al., 2006). This discrepancy may be attributed to mechanical forces in the rapidly proliferating post-implantation epiblast, which promote cell-cell contact and facilitate signaling required for gastrulation. Additionally, maternal factors present may compensate for mTORC2 loss during early development. Therefore, the role of RICTOR/mTORC2 in differentiation and development is multifaceted and influenced by both embryonic and environmental factors.

## Experimental procedures

### Resource availability

#### Corresponding author

Further information and requests for resources and reagents should be directed to and will be fulfilled by the lead contact,

#### Materials availability

The original hESC lines were purchased from WiCell under MTA, from which RIC-DF lines were generated. Thus, the RIC-RO hESC lines will be made available on request with completed Materials Transfer Agreements.

#### Data and code availability

The RNA-sequencing datasets of wildtype and RIC-KO hESCs are available in the GEO database with accession ID GSE243723.

### Culture and maintenance of hESCs

H1 (WA-01) and H7 (WA-07) hESCs from WiCell (https://www.wicell.org/) were routinely cultured on Matrigel-coated plates in hESC culture medium (MEF-CM with 10 ng/ml of bFGF (PeproTech)) with daily medium refreshment. MEF-CM was produced by incubating knockout serum replacement (KSR) medium (20% KSR, 1 mM L-glutamine, 1% non-essential amino acid, 0.1 M β-mercaptoethanol, 1% Penicillin-Streptomycin, 4 ng/ml bFGF in Knockout DMEM) on MEF for 24 hours (Noisa et al., 2011). hESCs were split at a 1:3 ratio by collagenase-assisted mechanical method. When required, hESCs were also dissociated into single cells by Accutase (Merck). All reagents were from Thermo Fisher Scientific unless stated. All hESC lines are negative for mycoplasma by regular test.

### Generation of *RICTOR*-knockout and RICTOR-knockdown hESCs

Targeting vector eSpCas9(1.1)-2A-Puro was constructed by replacing the Cas9 in pSpCas9(BB)-2A-Puro (PX459) with Cas9(1.1) from eSpCas9(1.1) (Addgene) (Ran et al., 2013; Slaymaker et al., 2016). Two sgRNAs (GGCCACAGTGAAGAAAAACTGGG and AGACTCCAGTATTCTCCAGA-AGG) were selected for co-transfection into H1 hESCs using the Neon^TM^ Transfection system (Thermo Fisher Scientific) following the manufacturer’s instruction (Figure S1A,B). Successful targeted colonies were selected through two-rounds of colony picking and screening. WT control clones were established in the same way but without sgRNA.

RICTOR-knockdown hESCs were established by transducing H7 hESCs with lentivirus expressing Rictor_2 shRNA (Addegene) as previously described (Sarbassov et al., 2005; Yu et al., 2015). Transduced cells were selected with puromycin (5 µg/ml) for a week from 48 hours post-infection.

### Differentiation via Embryoid body (EB) formation

EB formation was performed using AggreWell™ 400 plate (StemCell Technologies). The plate was prepared with anti-adherent rinsing solution and centrifuged at 13,00 g for 5 minutes. After removal of the solution, 7-8 x 10^5^ hESCs /well were seeded in EB medium (DMEM/F12 containing 20% fetal bovine serum or KSR, 1 mM Glutamine,1% Penicillin-Streptomycin, 1:100 N2 and 1:200 B27) with10 µM ROCKi and centrifugated at 100 g for 3 minutes. Cell aggregates were gently transferred to low attachment plates next day and cultured in suspension for another 6 days with medium change every 2 days. EBs were then plated to adherent culture dishes for further 7 days (Figure S2E).

### BMP4-induced mesendoderm differentiation

In colony: the method was adapted from previous ones (Bernardo et al., 2011; Warmflash et al., 2014). hESCs were dissociated into small clusters and seeded at a 1:5 ratio in hESC culture medium with 10 µM ROCKi that was withdrawn the next day. Differentiation started by adding 50 ng/ml BMP4 with or without 50 ng/ml AA or/and 6 μM CHIR into hESC culture medium for 48 hours (Figure S4E).

In spheroids: 5 x10^5^ hESCs/well were seeded into AggreWell™400 plate in hESC culture medium with 10 µM ROCKi and centrifuged at 100 g for 3 minutes. The medium was refreshed next day with 1 ng/ml BMP4 and 10 μM ROCKi and cultured for 48 hours (Figure S4D).

In single cells: hESCs were seeded on Matrigel-coated plates at 1 x 10^4^/cm^2^ in hESC culture medium with 10 μM ROCKi. Differentiation started the next day with fresh medium containing 50 ng/ml BMP4 for 48 hours (Figure 7A).

### Activin A-induced endoderm differentiation

hESCs were seeded on Matrigel-coated plates at 5 x 10^4^/cm^2^ in hESC culture medium with 10 μM ROCKi. Differentiation started by replacing hESC culture medium with RPMI1640/B27 medium containing 100 ng/ml Activin A for 72 hours with daily medium refreshing (Yu et al., 2015) (Figure S4C).

### Cell Adhesion assay

hESCs were seeded at 2.5 x 10^4^ cells/well on a Matrigel-coated 96-well plate in hESC culture medium containing 2.5-10 μM ROCKi and incubated at 37°C, 5% CO2 for 10 min, The attached cells were stained with 200 μl 0.2% crystal violet solution (Merck) for 10 min. After washing, 1% SDS (100 μl/well) was added and incubated for 15 min on a shaker and the data was recorded at 570 nm wavelength on an microplate reader (Bio-Rad).

### Alkaline phosphatase colony formation assay and CCK8 assay

hESCs were seeded at 6 x 10^3^ hESCs/well in 12-well plates in hESC culture medium for 5 days before stained with Leukocyte Alkaline Phosphatase Kit (Merck) following manufacturer’s instruction. Cell proliferation was measured using Cell Counting Kit-8 (CCK-8) from APExBIO following the manufacturer’s instructions and using a microplate reader.

### Reverse-Transcription-qPCR (RT-qPCR)

Total RNAs were isolated with TRI-Reagent® (Merck) and cDNA was synthesized with Superscript II reverse transcriptase (NEB). qPCR was performed in a StepOnePlus™ System (Applied BioSystems) using SYBR® Green JumpstartTM Taq Ready Mix (Merck). Table S4 lists all the primers.

### RNA-sequencing (RNA-seq) analysis

Total RNA was extracted using TRIzolTM Plus RNA Purification Kit (Thermo Fisher Scientific). Library preparation, sequencing, and initial quality check were performed by Imperial BRC Genomics Facility (IGF) using Illumina HiSeq 4000. Raw fastq data were trimmed by Fastp v0.23.2 (Chen et al., 2018) and STAR v2.7.10a (Dobin et al., 2013) was used to create genome mapping indices with human genome files (GRCh38 release 104). Transcripts were quantified by RSEM v1.3.3 (Li and Dewey 2011). All downstream analyses were carried out in R v4.1.2 using Bioconductor packages with variance differences and batch effects corrected (Ritchie et al., 2015). Expression analysis was performed by DESeq2 v1.34.0 (Love et al., 2014) with a false discovery rate (FDR) threshold of 0.05. Genes with a *p*-adj value < 0.05 and a log2 fold change (FC) >1 were considered differentially expressed. Gene ontology (GO) and Kyoto Encyclopedia of Genes and Genomes (KEGG) pathway analyses were performed separately on upregulated and downregulated genes using clusterProfiler v4.2.2 in R (Yu et al., 2012). *P*-values were adjusted using the Benjamini-Hochberg method with a cut-off at 0.05.

### Immunoblotting

Cell lysates were obtained in RIPA (Radio Immunoprecipitation Assay) buffer supplemented with phosphatase and protease inhibitors (all from Merck). 10-25 µg proteins were resolved in 8% SDS-PAGE gels and blotted onto polyvinylidene (PVDF) membranes, which were probed with primary antibodies overnight at 4°C followed by horseradish peroxidise (HRP) conjugated secondary antibodies for 1 hour before signals detected by ECL (enhanced chemiluminescent) and exposed onto CL-XPosure film (Thermo Fisher Scientific). Table S5 lists all the antibodies.

### Immunostaining and image analysis

hESCs cultured on Nunc Thermanox Coverslips (Thermo Fisher Scientific) or differentiated spheroids were fixed for 20 mins with 4% paraformaldehyde. After incubation with blocking buffer, cells were probed with primary antibodies overnight at 4°C, followed by 1 hour incubation with fluorescent conjugated secondary antibodies before counterstained with 4,6-diamidino-2-phenylindol (DAPI, Merck) and mounted onto microscope slides with Mowiol 4-88 solution. All the images, except those in Figure S4A, were captured with a Leica SP5 II confocal microscope. The images in Figure S4A were taken with Nikon Eclipse E400 microscope. Quantification of positive signals were performed using ImageJ and CellProfiler (McQuin et al., 2018) software after subtracting background signals using the Rolling Ball method with a radius of 50 pixels.

### RhoA activation assay and Rac1-GTP assay

RhoA activation assay was performed using RhoA G-LISA Activation Assay Kit – colorimetric format (Cytoskeleton Inc) following manufacturer’s instruction. The signals were recorded as absorbance at 490 nm wavelength using microplate spectrophotometer. Rac1-GTP assay was performed using Non-radioactive Rac1 Activation Assay Kit (Merck) following manufacturer’s instruction.

### Statistical analysis

All data are presented as the mean with error bars indicating standard deviation (SD), unless indicated otherwise, from at least three biological replicates; *p*-values were obtained by unpaired two-tail t test, one-way or two-way ANOVA using GraphPad Prism 9 except RNA-seq analysis.

## Acknowledgements

This research is supported by Genesis Research Trust (grant no. P64247, PA9387). FB is supported by PhD scholarship from the Punjab Educational Endowment Fund and YHC is partially sponsored by GSSA scholarship. We thank the Imperial BRC Genomics Facility (IGF) for providing the RNA-sequencing service and all the members in Stem Cell Differentiation group for their helpful discussion.

## Author contributions

W.C. conceived project and designed research strategy and experiments. Y-H.C. and F.B. designed gene targeting strategies and generated RIC-KO and -KD hESC lines, respectively and carried out initial characterization on them. L.T., F.B. and P.D.W. performed differentiation experiments and data analysis. L.T., F.B. and X.M. performed cell culture, molecular experiments and data analysis. L.T performed RNA-seq and analysis supervised by S.A. Y.Z. provided technical support for cell culture and molecular experiments. W.C. and L.T. wrote the manuscript with inputs from all authors.

