## Supplementary fig and fig legends for "mTORC2-mediated cell-cell interactions promote BMP4-induced WNT activation and mesoderm differentiation"

### SUPPLEMENTARY FIGURES AND FIGURE LEGENDS

Figure S1

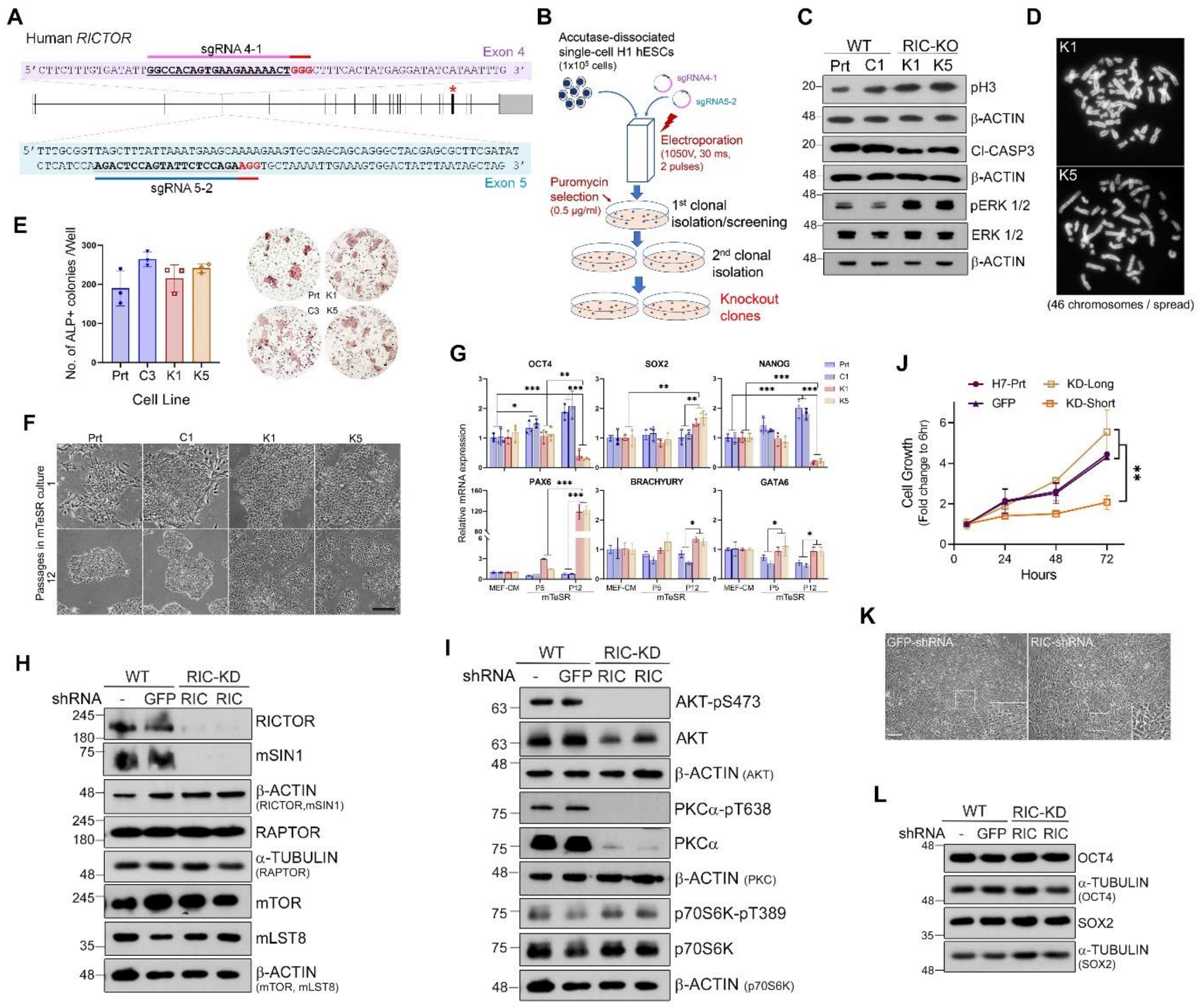


**Figure S1. Generation of RICTOR-knockout and -knockdown hESCs.**

(A) Diagram depicting human *RICTOR* gene, sgRNA targeting sequences and shRNA site (* in red).

(B) Schematic of generating *RICTOR*-KO clonal hESC lines.

(C) Representative immunoblots of the proliferation marker pH3 and apoptotic marker cleaved caspase 3 (Cl-CASP3) as well as ERK1/2-pThr202/Tyr204 in WT and RIC-KO hESCs (n = 3).

(D) Representative images of metaphase spreads of RIC-KO clones (n = 7 per line).

(E) Alkaline phosphatase colony formation assay. Quantitative analysis (left) shown as mean ± SD (n = 3) and representative images (right). Prt, C3 and K1, K5 represent H1 parental, control clone 3 and RIC-KO clone 1 and 5, respectively.

(F) Representative images of WT and RIC-KO hESCs in mTeSR cultures at passage 1 and 12. Scale bar = 200 µm).

(G)Comparison of mRNA expression in WT and RIC-KO hESCs under indicated culture conditions. Data are presented as mean ± SD (n = 3).

(H) Immunoblots of RICTOR and mTOR complexes protein subunits in RICTOR-knockdown (RIC-KD) H7 hESCs.

(I) Immunoblots show abolished mTORC2 activities in RIC-KD H7 hESCs.

(J) Proliferation of WT and RIC-KD H7 hESCs by CCK8 assay. Indicated passage numbers of H7 RIC-KD are counted after selection. Data presented as mean ± SEM (n = 3).

(K) Representative phase-contrast images of H7 WT and RIC-KD hESCs. Scale bar = 100 µm.

(J) Immunoblots of key pluripotent transcription factors in WT and RIC-KD hESCs. All the immunoblotting experiments are repeated at least once with different culture passages (n ≥ 2).

*, ** & ***, *p* <0.05, 0.005 & 0.0005, respectively by two-way ANOVA.

Figure S2

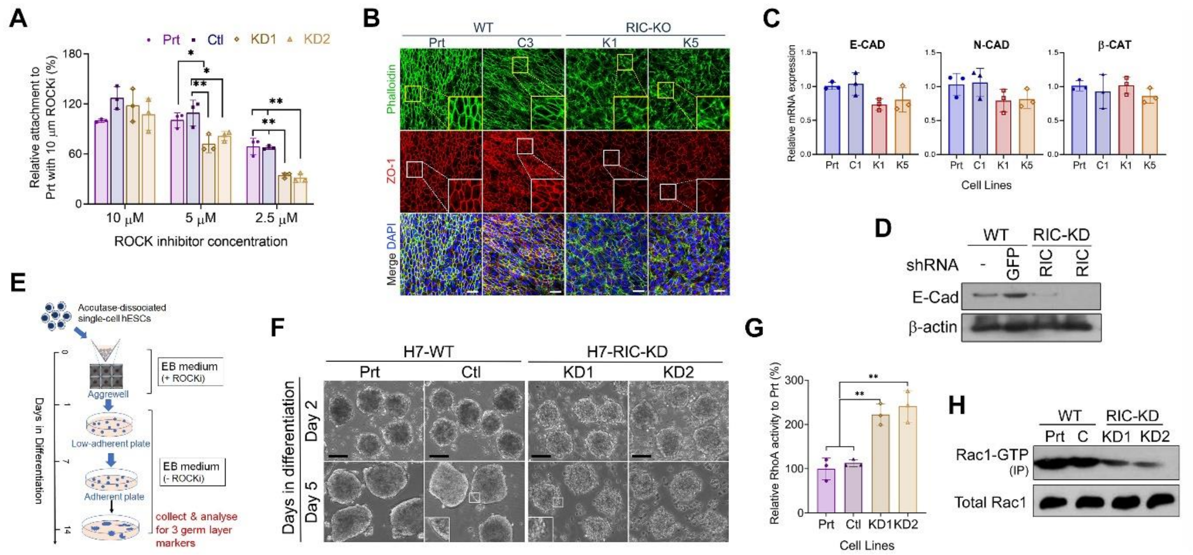


**Figure S2. Effects on cell adhesion in RIC-KO and RIC-KD hESCs.**

(A) Cell attachment assay in RIC-KD H7 hESCs as described in Figure 2B.

(B) Expression of ZO1 and F-ACTIN (Phalloidin) in H1 RIC-KO hESCs by immunostaining. Scale bar = 50 µm. For the enlarged inserts, the scale bar = 25 µm.

(C) RT-qPCR showing expression of adhesion complex genes, β-CATENIN (β-CAT), E- and N-CADHERIN (E-CAD and N-CAD) in H1 RIC-KO hESCs. Data are presented as mean ± SD (n = 3).

(D) E-CADHERIN expression in H7 RIC-KD and WT hESCs by immunoblotting (n = 2).

(E) Schematic illustrating the procedure of EB formation.

(F) Representative images of H7 RIC-KD EB formation in the presence of ROCKi. Insert showing blebs in RIC-KD EBs. Scale bar = 100 µm.

(G) RhoA activity in H7 WT and RIC-KD hESCs. Data are presented as mean ± SD (n = 3). ** *p* < 0.005 by one-way ANOVA.

(H) Rac1-GTP in WT and RIC-KD H7 hESCs (n = 2). KD1 & KD2 are derived from two knockdown experiments.

Figure S3

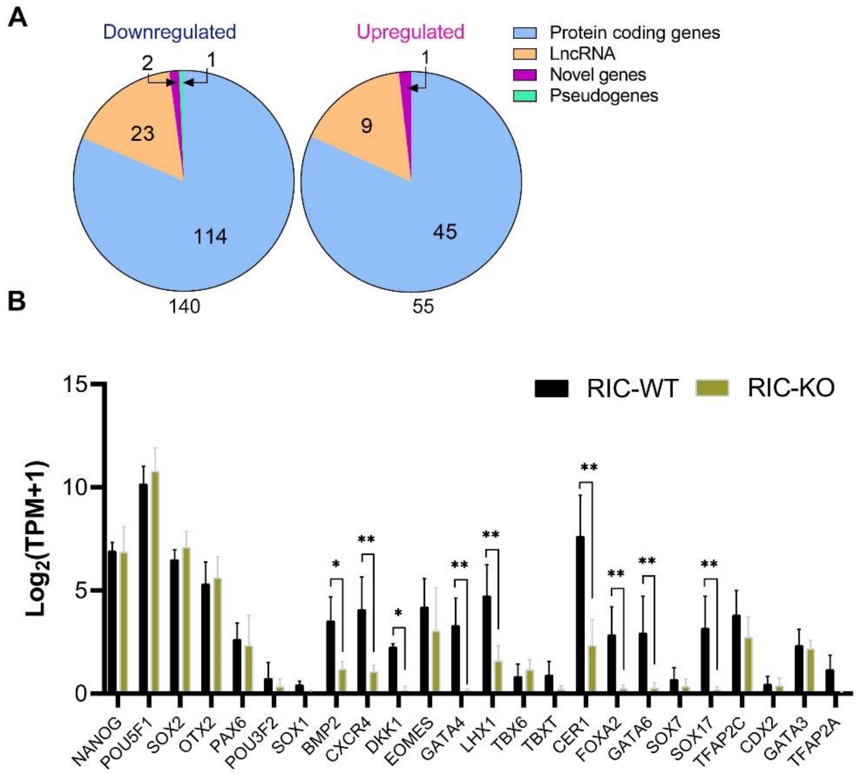


**Figure S3. Differentially expressed genes in RIC-KO hESCs by RNA-seq analysis.**

(A) Pie-chart showing distribution of differentially expressed genes in RIC-KO hESCs.

(B) Bar chart comparing the expression of representative lineage-associated transcription factors between WT and RIC-KO hESCs. Data are presented as mean ± SD (n = 4). * & **, $p < 0.05$ and 0.01, respectively by unpaired two tail t test.

Figure S4

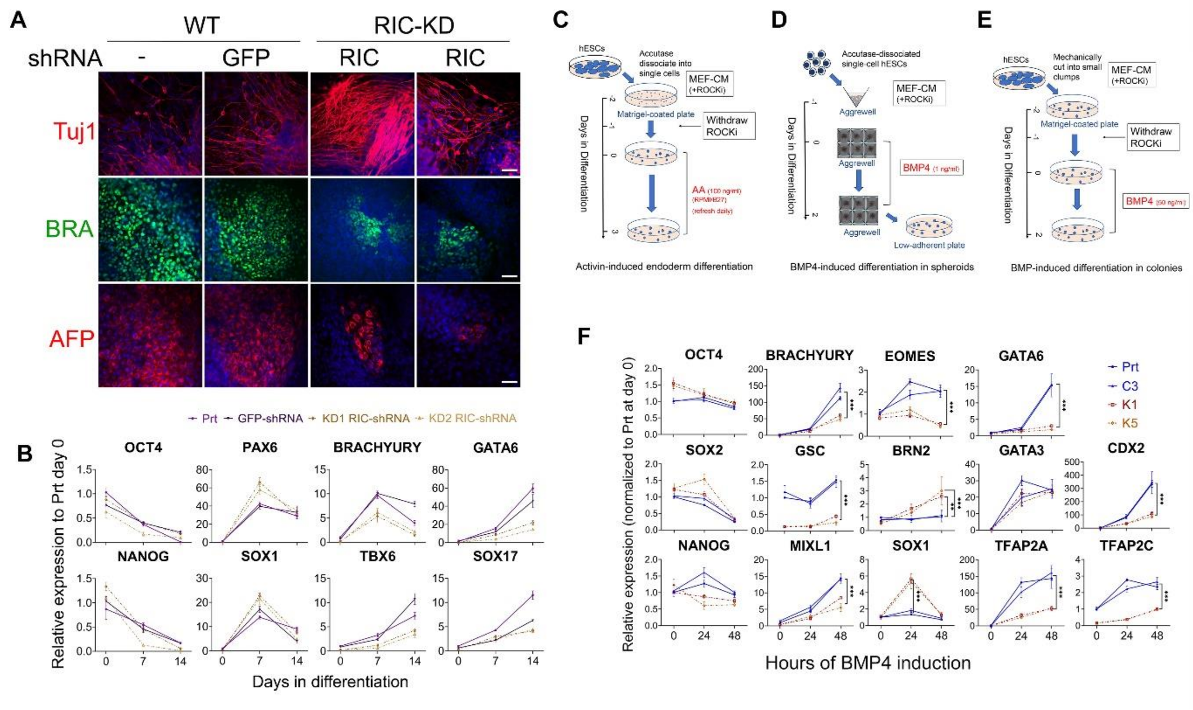


**Figure S4. Reduced mesoderm/endoderm differentiation in RIC-KD and RIC-KO hESCs.**

(A) Immunostaining of the three germ layer markers in WT and RIC-KD H7 hESCs after 14 days of EB differentiation (n = 2). Scale bar = 50 µm. The RIC-KD images represent two independent KD lines.

(B) Dynamic expression of indicated genes during EB differentiation in WT and RIC-KD H7 hESCs. Data are presented as mean ± SD of 6 measurements of 2 independent differentiation experiments.

(C) Schematic of Activin-induced endoderm differentiation.

(D) Schematic of BMP4-induced differentiation in spheroids.

(E) Schematic of BMP4-induced differentiation in hESC colonies.

(F) Expression of various lineage markers by RT-qPCR in BMP4-induced differentiation in WT and RIC-KO hESCs. Data are presented as mean ± SD (n = 3). ** & ***, *p* <0.005 and 0.0005 by two-way ANOVA.

Figure S5

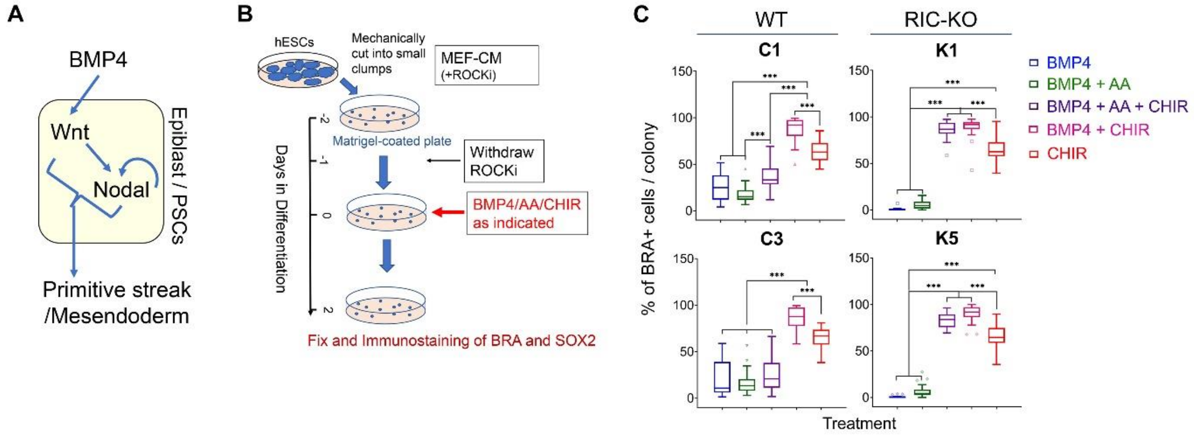


**Figure S5. WNT and Activin signaling differentially affect BMP4-induced mesendoderm differentiation in WT and RIC-KO hESCs.**

(A) Diagram illustrating signaling pathways involved in the regulation of gastrulation and mesendoderm differentiation.

(B) Schematic of the experimental procedure in the differentiation of hESCs with combined activation of BMP4, WNT and AA (Activin A) pathways.

(C) Comparing the effect of indicated pathways on differentiation of BRA+ cells within the same cell line. Data are presented as the Tukey box plot with number of colonies shown in Figure 4F and Figure 5 (n = 3). *, ** and ***, $p < 0.05$, 0.005 and 0.0005, respectively, by one-way ANOVA.

Figure S6

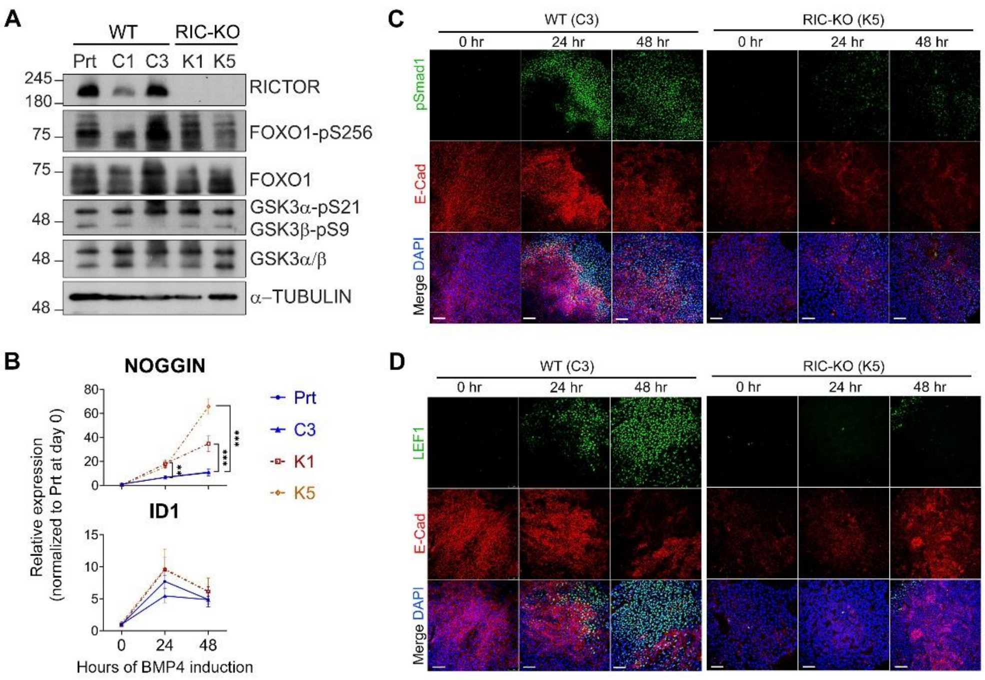


**Figure S6. E-CADHERIN modulates BMP4-induced pSMAD1 signal and LEF1 expression.**

(A) GSK3 and FOXO1 phosphorylation show similar patterns in WT and RIC-KO hESCs by immunoblotting (n = 2).

(B) Upregulation of BMP4 signaling target genes by RT-qPCR. Data are presented as mean ± SD(n = 3). ** and ***, $p < 0.005$ and 0.0005, respectively by two-way ANOVA.

(C,D) Immunostaining with indicated antibodies in C3 and K5 hESCs as described in Figure 6C,D (n = 3). Scale bar = 100 µm.

Figure S7

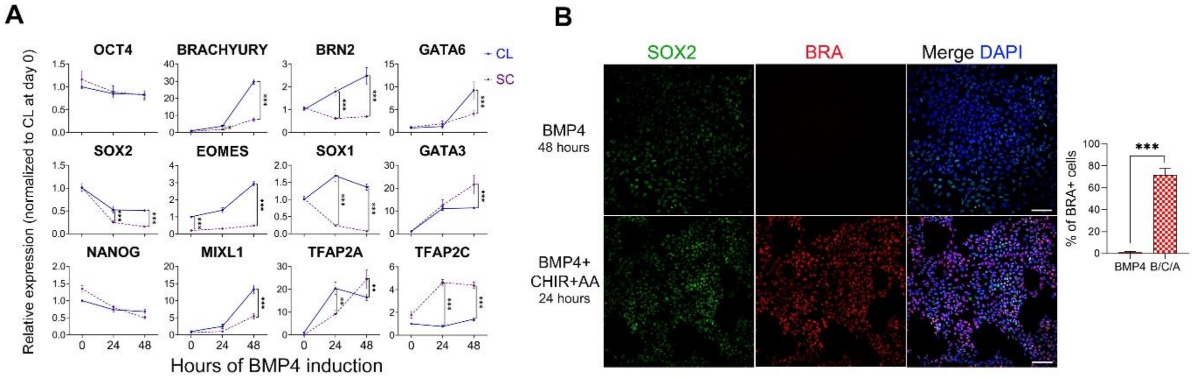


**Figure S7. Cell-cell contacts affect BMP4-induced differentiation.**

(A) RT-qPCR showing dynamic expression of indicated pluripotent and lineage genes during BMP4-induced differentiation in WT hESCs seeded in colonies (CL) or single cells (SC). Data are presented as mean ± SD (n = 3). ** & *** represent $p$ < 0.005 and 0.0005 respectively by two-way ANOVA

(B) Immunostaining of BRACHYURY (BRA) and SOX2 after differentiation of SC-seeded WT hESCs by indicated treatment. Representative images are shown (left) with quantification of BRA+ cells (right) with data presented as mean ± SD of 3 independent differentiation experiments with at least three random-selected fields being counted for each experiment. ***, $p$ <0.0005 by unpaired t-test. CHIR, CHIR99021; AA, activin A. Scale bar = 100 µm.
